# A delivered DNase toxin creates population heterogeneity through transient intoxication of siblings

**DOI:** 10.1101/2024.12.20.629666

**Authors:** Hanna Eriksson, Susan Schlegel, Jonas Kjellin, Sanna Koskiniemi

## Abstract

Population heterogeneity is important for multicellular behavior and division of labor. Bacterial toxin delivery has been implicated in generating population heterogeneity, but the molecular mechanisms behind this are not well understood. Here we investigate <u>how</u> CdiA toxins generate heterogeneity in isogenic populations. Using a DNase toxin as proxy, we find that *E. coli* populations able to deliver the toxin show a heterogeneous expression of the SOS-response gene *sulA*. Heterogeneity results from excessive delivery of toxin into some cells, which become intoxicated due to insufficient immunity. Intoxication is transiently reversible, and intoxicated cells can be rescued by *de novo* synthesis of cognate immunity protein. Expression of *sulA* is regulated by both DNA damage and redox status. Interestingly, kin-delivery changes redox status, whereas intoxicated non-kin cells induce the SOS DNA damage response. The former results in changed expression of metabolic genes whereas the latter induces prophage excision, which may promote horizontal gene transfer. In conclusion, we identify a molecular mechanism by which heterogeneity is generated through toxin delivery among kin, and the consequences of said heterogeneity.

**Significance statement:** Bacteria communicate through secretion of chemical signaling molecules to perform multicellular behavior. Recent advances suggest that contact-mediated toxin delivery allow bacteria to participate also in direct cell-cell communication. How such toxin-mediated communication would work mechanistically is however unclear. Here we elucidate a molecular mechanism of a toxin-mediated communication, where kin-cells transiently intoxicate each other, resulting in physiological changes. These changes depend on the toxic activity, i.e. other toxins with different activities are likely to give rise to other responses. Thus, the arsenal of toxins that a bacterium harbors could affect their ability to communicate. Understanding the molecular mechanism of how toxins could mediate polyphenism is important for our understanding of what this signaling is used for.

## Main text

Most bacterial species participate in multicellular behavior such as biofilm formation [1], for which communication is essential. To communicate, bacteria secrete soluble chemicals that allow them to sense the presence of other bacteria of the same species, a phenomenon known as quorum sensing [2]. In “true” multicellular organisms, direct cell-to-cell signaling is found alongside the production of soluble signaling molecules [3]. Early work suggested that contact-dependent growth inhibition (CDI), a toxin delivery system in bacteria, could be used for contact-dependent signaling (CDS) during biofilm formation [4]. Although recent findings show that toxin delivery is not relevant in this context [5], accumulating evidence suggests that interbacterial toxin delivery through contact-dependent mechanisms, results in population heterogeneity and changes in gene expression [6, 7]. This suggests that toxin delivery systems could function in inter-bacterial signaling. However, how bacterial toxins mediate population heterogeneity on a molecular level is still unclear, as are potential benefits of such heterogeneity.

Toxin-mediated heterogeneity in gene expression could arise by at least two distinct molecular mechanisms: i) the toxins, either by themselves or in concert with the immunity protein, could act as transcriptional regulators, as has been seen with type II TA-systems [8, 9] or ii) the toxins could change gene expression through their toxicity, as seen for e.g. type II toxins that specifically cleave certain mRNA [10] . Both toxin and immunity of TA-systems are expressed inside the cell, and if and when the toxin:antitoxin balance is disrupted is still unclear for many systems. Delivered toxins on the other hand possess a built-in ability to change the ratio between toxin and antitoxin as kin-delivery of a toxin will increase the number of toxins in the recipient cell, changing the toxin:antitoxin ratio and possibly affecting gene-expression.

In *E. coli*, toxin delivery can be mediated through <u>c</u>ontact <u>d</u>ependent growth inhibition (CDI) [11]. Cells with CDI systems express a stick-like CdiA protein on their cell surface. Upon contact with a cell expressing the cognate outer-membrane receptor (e.g., BamA) CdiA is cleaved at a conserved VENN motif, resulting in transfer of the <u>C</u>-terminal toxin domain (CdiA-<u>CT</u>) into the target cell periplasm (Figure 1A) [12]. To access the cytosol or inner membrane, where characterized CdiA toxins exert their function, the N-terminal part of the CT (also known as the <u>C</u>-terminal <u>e</u>ntry <u>d</u>omain, or CED) interacts with a cognate protein in the inner membrane that facilitates cytosol or membrane entry in an as yet undefined manner [13]. CdiA-CTs are polymorphic toxins with different biological activities, including ionophores and nucleases with DNase or RNase activity [14, 15]. To avoid self-inhibition, cells produce as immunity protein (CdiI) that protects against its corresponding CdiA-CT toxin [16].

**Figure 1.**
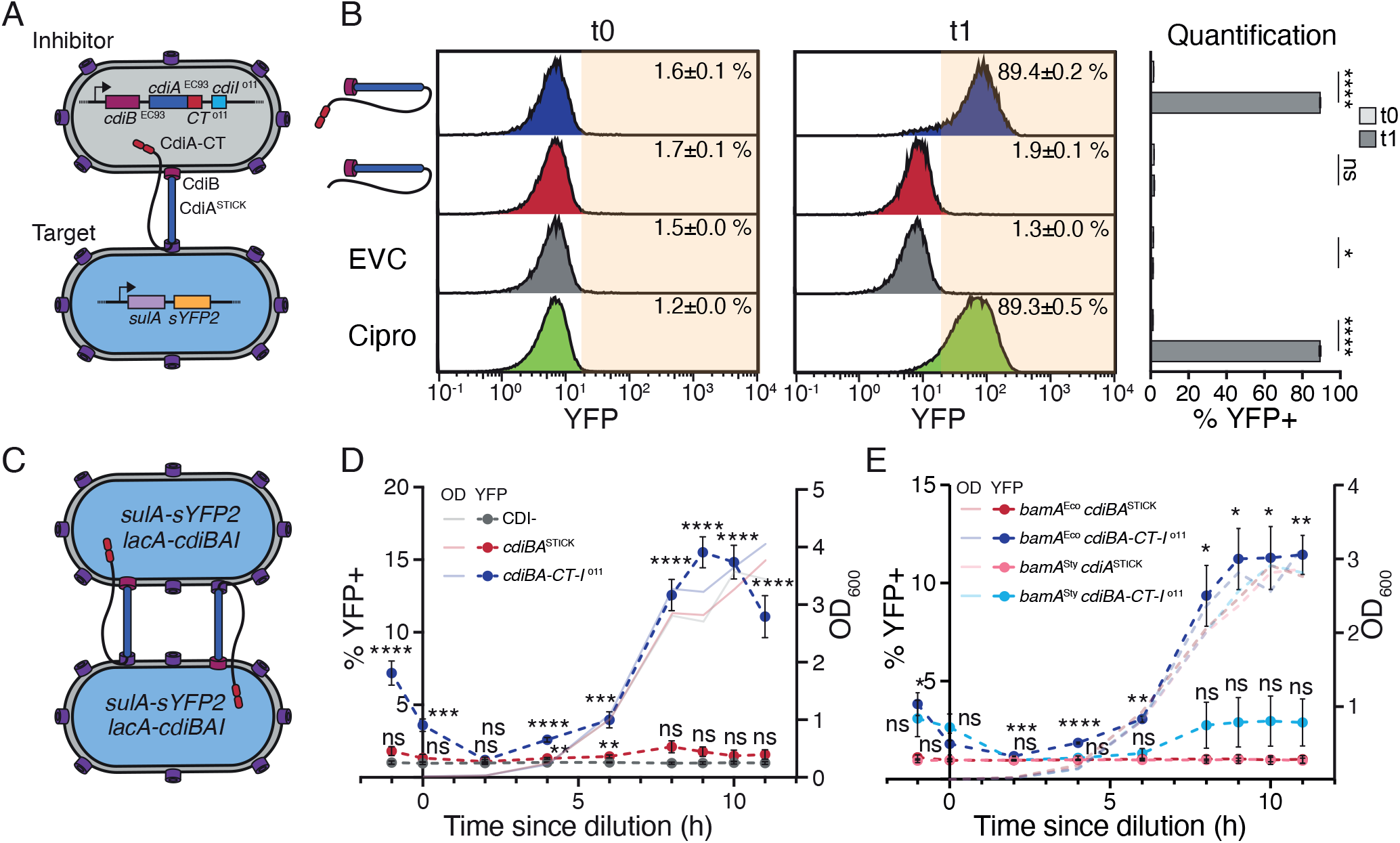
CDI+ *E. coli* intoxicate their siblings. **A)** Overview of strains used for competition in B. **B)** Competition between MG1655 inhibitors with or without p*cdiBA-CT-I* ^o11^ and MG1655 targets with *sulA*-*sYFP2* reporter. As a positive control for *sulA-sYFP2* induction, 0.1 mg/L ciprofloxacin was added to target cells in the absence of inhibitor. Enumeration of YFP+/-cells through flow cytometry as shown in histograms and in bar chart. t0 = before mixing, t1 = after 1 h of co-culture. EVC = empty vector control. (N= 6, biological replicates). **C)** Schematic of CDI+ strain used for monoculture experiment in D. **D-E)** Time-resolved enumeration of YFP+ cells in monocultures grown in M9-gly-CAA for 11 h. MG1655 with p*cdiBA-CT-I* ^o11^ (blue) or p*cdiBA*^STICK^ (red) were used in D and MG1655 with p*cdiBA-CT-I* ^o11^ or p*cdiBA*^STICK^ either with cognate *bamA*^Eco^ (red and blue) or non-cognate *bamA*^Sty^ (pink and light blue) were used in E. CDI-(grey) = MG1655 with the *sulA*-*sYFP2* reporter. (N= 12 (D) or N=4 (E), biological replicates). Error bars are SEM. Statistical significance was determined through B) Student’s t-test and D-E) two-way ANOVA with Tukey’s posthoc test. * <0.05, ** <0.01, *** <0.001, ****<0.0001. Significance in D, E is relative to the respective *cdiBA*^STICK^ strain.

Here we set out to investigate if and by what mechanism a CdiA toxin causes population heterogeneity, using a toxin with DNase activity as a proxy. We find that a subpopulation of cells expressing this CDI system becomes intoxicated during dense growth in minimal M9 media. Intoxication is delivery-dependent and appears to occur when the toxin molecules outnumber the antitoxins, rather than being a consequence of antitoxin instability. Intoxicated cells can resume growth after *de novo* synthesis of immunity protein to a certain level of intoxication, indicating a reversible process. These cells change their gene-expression, but the changes depend on the degree of intoxication. During kin-delivery, changes in gene-expression appear related to altered redox status in response to the toxin. In contrast, cells intoxicated by non-kin, induce the SOS DNA damage response, resulting in cell death and prophage excision. Taken together, our results suggest that kin-delivery of toxins could function as a contact-dependent signaling mechanism by which the redox status and gene-expression pattern of a subpopulation of cells is changed at high population densities. This is in contrast to previously identified roles of toxin delivery among non-kin cells.

## Results

### Delivery of CdiA-CT^o11^ toxin among kin induces *sulA* expression in a subpopulation

To investigate if kin-delivery of a DNase toxin could generate population heterogeneity, we used a previously described yellow fluorescent protein (YFP) based transcriptional reporter, *sulA-sYFP2*, that signals when toxicity occurs in an individual cell. DNA damage induces the LexA-controlled SOS DNA damage response genes in *E. coli* including the cell-division inhibitor SulA [17]. To investigate if delivery of the CdiA-CT^o11^ toxin (with known DNase activity [15]) induced *sulA-sYFP2*, we competed E.coli with plasmid-encoded p*cdiBA-CT-I*^o11^ locus against target cells with the reporter. Targets constitutively expressed blue fluorescent protein (BFP) [11] and inhibitors red fluorescent protein (dTomato) to enable separation of the two (Fig. 1A). Using flow cytometry, we found that 89% of the target cells became YFP-fluorescent after 1 h of co-culture with inhibitors, indistinguishable from the ciprofloxacin (CIP)-treated positive control (Fig. 1B). In contrast, less than 2% YFP+ cells were observed when inhibitor cells carried either an empty vector control devoid of CDI (EVC) or an empty stick control (p*cdiBA*^STICK^) lacking *cdiA-CT-I* ^o11^ (Fig. 1B). In agreement with the fluorescence data, no competitive advantage (monitored by viable counts on selective agar plates and summarized as competitive index) was observed for inhibitor control strains (EVC, p*cdiBA*^STICK^), while a 100-fold advantage was observed for inhibitors carrying the CdiA-CT^o11^ toxin (Fig. S1A). Thus, the *sulA-syfp2* reporter is a functional readout for CdiA-CT^o11^ intoxication.

As the majority of CDI systems are found in single copy on the bacterial chromosome [18] we incorporated the *cdiBA-CT-I* ^o11^ or the *cdiBA*^STICK^ locus into the chromosome. Cells carrying chromosomal *cdiBA-CT-I* ^o11^ induced an SOS response in around 77% of sensitive target cells and inhibited growth (Fig. S1BC), confirming functional toxin delivery. To investigate whether the presence of CdiA-CT^o11^ can cause population heterogeneity, *sulA-sYFP2* positive cells with either *cdiBA-CT-I* ^o11^ or *cdiBA*^STICK^ on the chromosome (Fig. 1C) were grown in monoculture and the fraction of YFP+ cells in the population over time was monitored using flow cytometry. At higher cell densities (OD_600_ ≈3, Fig. S1D), *sulA*-*sYFP2* was induced in >10% of the cells expressing *cdiBA-CT-I* ^o11^ while no increase in fluorescence was observed for either the *cdiBA*^STICK^ or the non-CDI control (Fig. 1D). To confirm that *sulA*-*sYFP2* induction was indeed due to delivery of toxin, the experiment was repeated using a strain in which delivery was restricted by replacing the native, cognate receptor *bamA*^Eco^ by a non-cognate one from *Salmonella enterica* serovar Typhimurium LT2, *bamA*^Sty^ [19]. In these monocultures, no increase in the fraction of YFP+ cells was detected (Fig. 1E), suggesting that kin-delivery of CDI can cause heterogeneous *sulA* expression in the population.

### Intoxication of cells by increasing delivery of toxin

The intoxication of only a subpopulation of cells raises questions regarding the molecular mechanism at play, i.e. what makes some cells in an isogenic population sense the toxin while their siblings do not? A reasonable assumption is that the toxins outnumber the immunity proteins in the affected cells.

To investigate whether variations in the toxin:immunity ratio could arise from differences in toxin or immunity protein turnover rates, the stability of CdiA-CT^o11^ and CdiI^o11^ was assessed by western blot following DL-serine hydroxamate (SHX)-mediated translation arrest. Whereas CdiI^o11^ appears completely stable even 120 minutes after translation arrest (Fig. S2AB), CdiA-CT^o11^ with a C-terminal alfa-tag (CdiA-CT^o11^-α) was degraded below the detection limit at the same time point (Fig. S2C). This suggests that the observed heterogeneity is unlikely to be due to proteolytic turnover of the immunity protein.

To assess if kin-intoxication instead could originate from an over-delivery of toxin relative to the protective capacity of individual cells, we tested whether cells with immunity become intoxicated when too much toxin is delivered to them. To control the level of intoxication, target cells were supplemented with a plasmid expressing arabinose-inducible *cdiI*^o11^-α (p*cdiI*^o11^-α). The level of CdiI^o11^-α in the population was altered using a method inspired by the fluorescence dilution assay [20]. Exponentially growing target cells were either induced (“+”) with 0.02 % L-arabinose for 15 minutes to obtain a homogeneous expression of *cdiI*^o11^-α in individual cells (Fig. S3A) or left uninduced (“-”). Subsequently, cultures were diluted and re-grown in 8 cycles of 2 h each, equating to ∼19 generations (Fig. 2A). Assuming that degradation is negligible (Fig. S2AB), the amount of CdiI^o11^-α should decrease by half at every cell division, resulting in a ∼5-fold dilution during each 2 h cycle (Fig. S3B, right panel). At the end of each cycle, samples were taken for i) a competition against an inhibitor strain with p*cdiBA-CT*-α *-I* ^o11^(Fig. 2B) and ii) western blotting to monitor levels of immunity and toxin (Fig. 2CD). As expected, the level of toxin produced by the inhibitor population remained roughly constant over time (Fig. 2D). In contrast, immunity levels decreased with the anticipated ∼5-fold dilution per cycle after cycle 2 (Fig. 2C and S3B), indicating that any residual immunity induction had ceased. The induced samples also showed decreasing protection over the course of the experiment (blue bars, Fig. 2B). At cycle 5, target cells were outcompeted 10-fold (Fig. 2B), indicating that the levels of CdiI^o11^-α were too low to protect from delivered toxin in some cells. At later time points, the fraction of CDI-sensitive cells equaled that of non-induced control cells (red bars, Fig. 2B). To exclude accumulation of resistance mutations, target cultures were split at cycle 8 and expression of immunity was re-induced in ½ of each culture irrespective of previous induction status (cultures termed 8+). Cells from both cultures were again fully protected against CdiA-CT^o11^ and showed full immunity expression (Fig. 2BC). Taken together, these results suggest that growth is inhibited when toxin delivery exceeds the capacity of the available immunity proteins.

**Figure 2.**
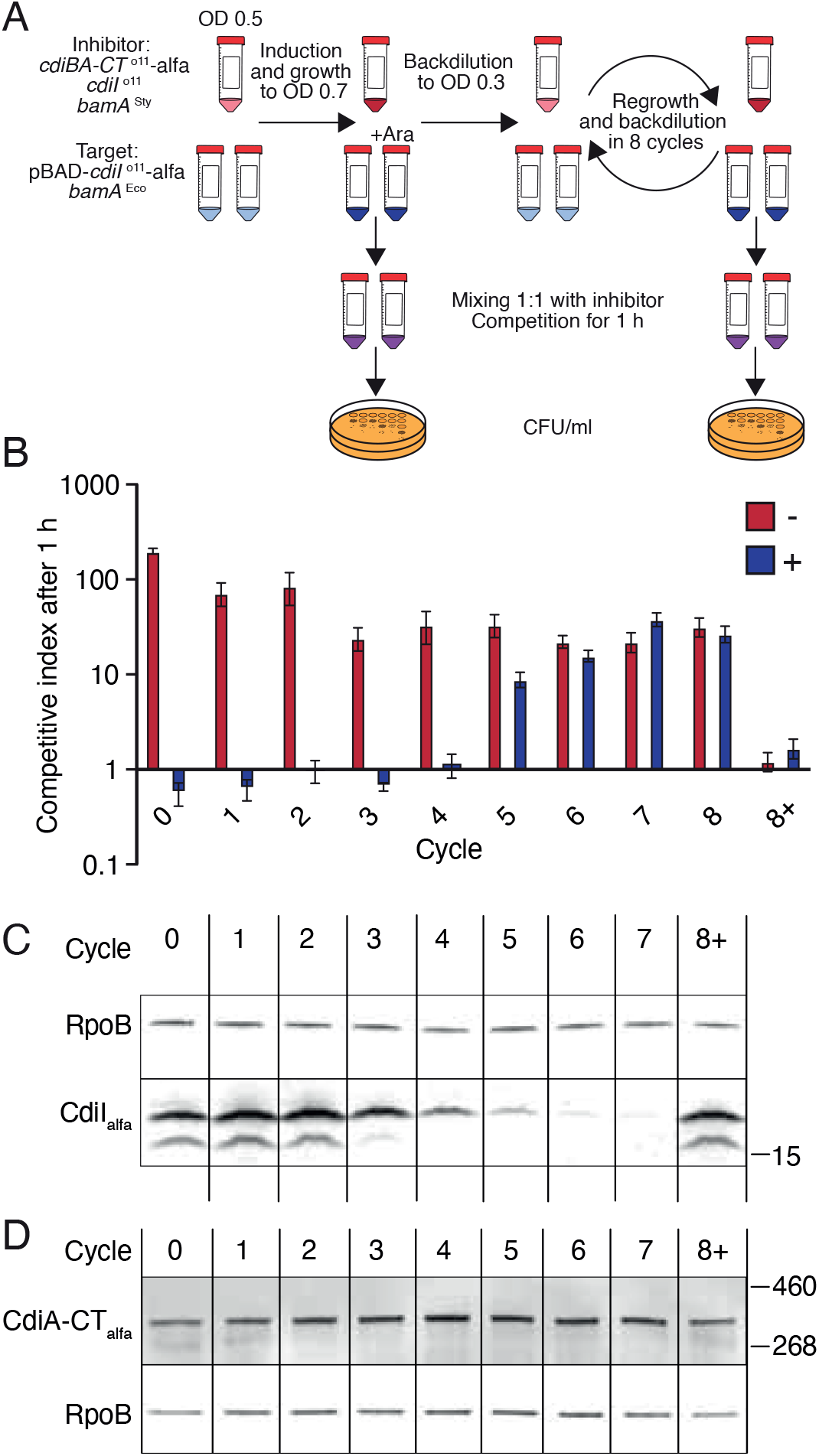
Target cells become intoxicated when toxin levels outnumber immunity levels. **A)** Overview of the experiment. **B)** Competitive index after co-culturing p*cdiBA-CT-*α*-I* ^o11^ inhibitors with targets containing different levels of CdiI^o11^-α immunity (as seen in C) for 1h. Red /blue bars indicate target cells grown in the absence /presence of L-arabinose before cycle 0. Competitions were performed after each cycle. At cycle 8 the culture was split and half of the culture was supplemented with L-arabinose (8+) to verify that immunity induction was still possible. (N = 3 biological replicates). Error bars are SEM. **C)** Western blot to detect CdiI^o11^-α in target cells using anti-α nanobodies. **D)** Western blot to detect the level of full-length CdiA^o11^-α in inhibitor cells (expressed by the inhibitor throughout the different cycles of the experiment) using anti-α nanobodies.

### Intoxication by CdiA-CT^o11^ is reversible within a short time window

For an intoxication event to result in a response other than growth inhibition, irrespective of the underlying mechanism, the damage caused by the toxin must be reversible. To investigate whether CdiA-CT^o11^-intoxicated cells can be rescued, cells with p*cdiBA-CT-I*^o11^ were competed against target strains carrying either an arabinose-inducible *cdiI* plasmid (IndImm) or an empty vector control (NC). Arabinose was added to the cultures 0, 5, 10, 30 or 60 minutes after mixing (Fig. 3A). The cells were left to recover for 30 min before fluorescence was assayed, whereas viability was assessed immediately. After 5 minutes, only 25% of the non-immune target cells (NC) could form colonies (Fig. 3B, left panel), and 20% induced *sulA-sYFP2* expression (Fig. 3B, right panel). This suggests that excessive damage prevents some of the intoxicated cells from inducing the DNA damage response. When immunity was induced after 5 min of competition, 50% of the targets formed colonies, and 34% were YFP+ (Fig. 3B). After 10 minutes of competition, only 5-to 10% of either target formed colonies and YFP+ cells increased to ∼50% (Fig. 3B).

**Figure 3.**
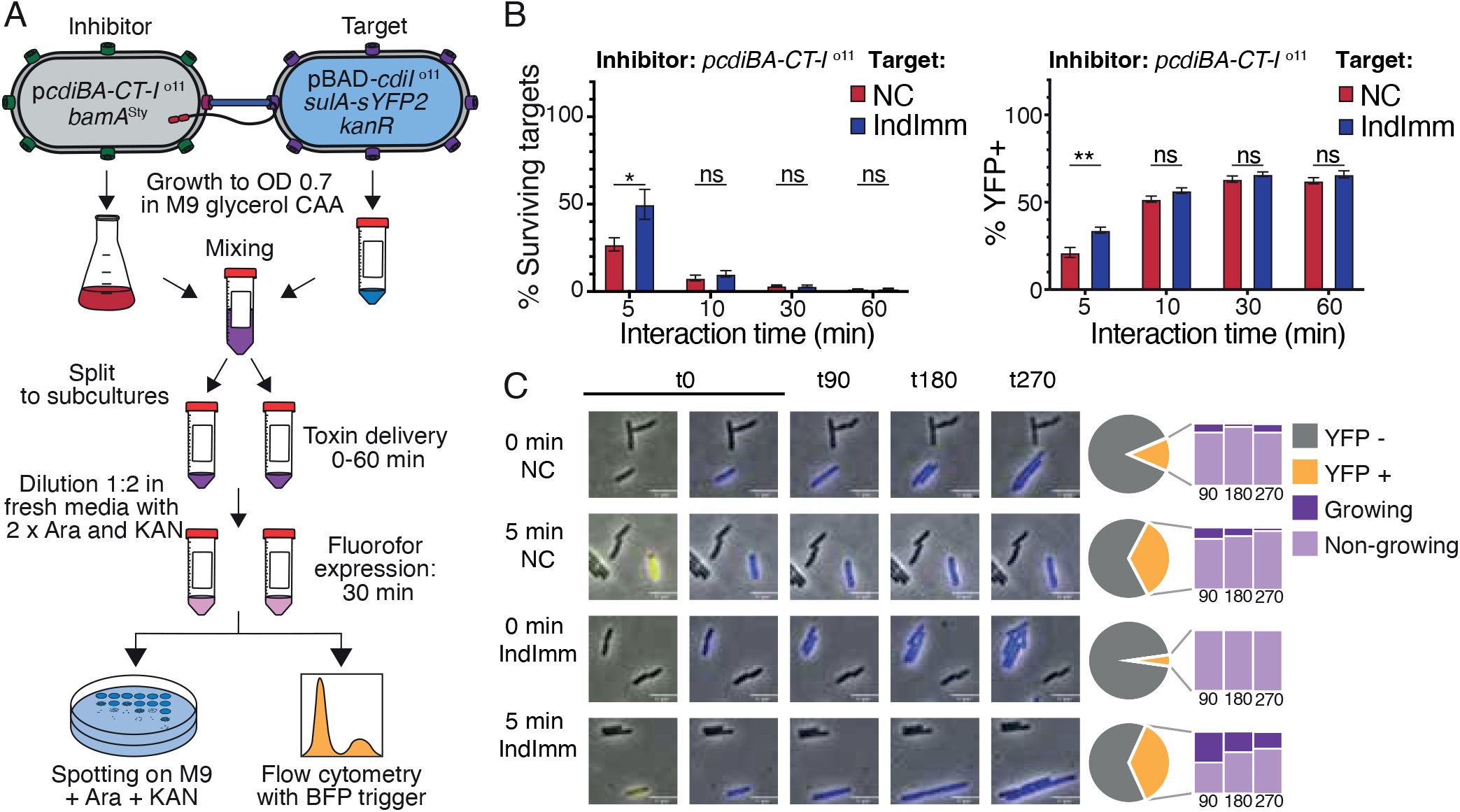
Cells intoxicated by CdiA-CT^o11^ resume growth upon induction of *cdiI*^o11^. **A)** Overview of the experimental setup for B. Target cells are KanR and BFP+ to make them distinguishable from inhibitors. **B)** % surviving (compared to t0) and % YFP+ (compared to BFP+ cells at the given time point) target cells with (IndImm, blue bars) or without (NC, red bars) pBAD::*cdiI* ^o11^ after co-culture with p*cdiBA-CT-I* ^o11^ inhibitors for 5, 10, 30 or 60 minutes. (N = 8 biological replicates). Error bars are SEM. Statistical significance was determined using Students t-test. * <0.05, ** <0.01. **C)** Time-lapse microscopy on cells from the competition in B. Cells are assessed as YFP+ (yellow) or YFP-(grey). Fraction of YFP+ target cells that do (dark purple) or do not (light purple) resume growth after 90, 180 and 270 minutes of recovery is shown.

Even though these results indicate that intoxication inflicted by a DNase toxin can indeed be reversed – if only for a short time - bulk experiments do not reveal if the observed growth occurs in the cells that experience DNA damage or in the non-affected population. We therefore repeated the assay described in Figure 3A but conducted the recovery phase on agarose pads and followed growth resumption of individual cells by time-lapse microscopy. Both p*cdiI* ^o11^ and non-immune cells did resume growth when recovery was initiated immediately after mixing with inhibitor cells (0 min). Only few cells showed signs of *sulA-sYFP2* expression (Fig. 3C and S4A), indicating that negligible amounts of toxin had been delivered. After 5 minutes of competition, the fraction of YFP+ cells had increased similarly in both target strains, but after 90 minutes of recovery the fraction of YFP+ targets that could resume growth was higher in p*cdiI* ^o11^ cells (50%) as compared to the empty vector control (18%) (Fig. 3C and S4BC). At later timepoints the number of cells that resumed growth decreased in both conditions, likely because some cells lyse after resuming growth. Even at these timepoints the fraction that could resume growth was larger in p*cdiI* ^o11^ cells than in the control (Fig.3C). These results show that increasing levels of immunity can mitigate the effects of intoxication by CdiA-CT^o11^in part of the population.

### CdiA intoxication changes cellular metabolism and redox status

Our experiments demonstrate that kin-intoxication by CdiA toxins can induce population heterogeneity in terms of gene expression, but do not shed light on the evolutionary purpose of such heterogeneity. We reasoned that a more comprehensive understanding of what happens in CdiA-CT^o11^-intoxicated cells, could provide valuable leads. Therefore, we cloned CdiA-CT^o11^ under an arabinose-inducible promoter in cells with a chromosomal *cdiBA-CT-I*^o11^ locus (to mitigate toxicity from leaky expression). 0, 5 and 20 minutes after the addition of arabinose, mRNA levels were monitored using transcriptomics. Differential expression was determined relative to an EVC strain (without the *cdiBAI* operon on the chromosome) at each time point. Expression of CdiA-CT^o11^ reduced viability of the cells, with 10% survival after 5 minutes of induction and 0.1% survival after 20 minutes of induction (Fig. S5A). 5 minutes after CdiA-CT^o11^ induction we observed a moderate increase in mRNA levels for a small subset of genes involved in, e.g. the SOS-response (*sulA, recN*), phenylacetyl-metabolism (*paaA-H*), and prophages (*exoD, ybcVW, nohB*). Similarly, only few loci showed a decrease in mRNA levels, e.g. *rlmG, ygiQ, opgC, tdcEFG* and *fdoGHIE*. Keeping in mind that the average half-life of mRNAs in *E. coli* is estimated to be 3-8 minutes [21] it is not surprising that we did not see a stronger decrease at this time-point. Genes affected by either the addition of Arabinose (*araBAD, araE, araC, ygeA*) or the chromosomal *cdiBAI-*locus (*lacA*) were not considered further.

After 20 minutes, more than 2000 genes showed differential mRNA levels compared to the empty vector control (Fig. 4A). To assess which of the time-points best represents the physiological state of the YFP-positive cells in the monoculture (compare Fig. 1), we used a previously described reporter for the DNA damage inducible gene *tisB* [17]. At 20 minutes *tisB*-mRNA levels were altered similarly to *sulA* (42- and 35-fold increase, respectively), but in contrast to *sulA*, no increase in mRNA levels could be observed at 5 minutes (Fig. S5B). Despite being induced by the delivery of CdiA-CT^o11^ in a competition (Figure S5CD), no *tisB*-*sYFP2* positive cells could be observed in monoculture, suggesting that the 5-minute time point best represents the level of intoxication observed in a monoculture (Fig. S5E).

**Figure 4.**
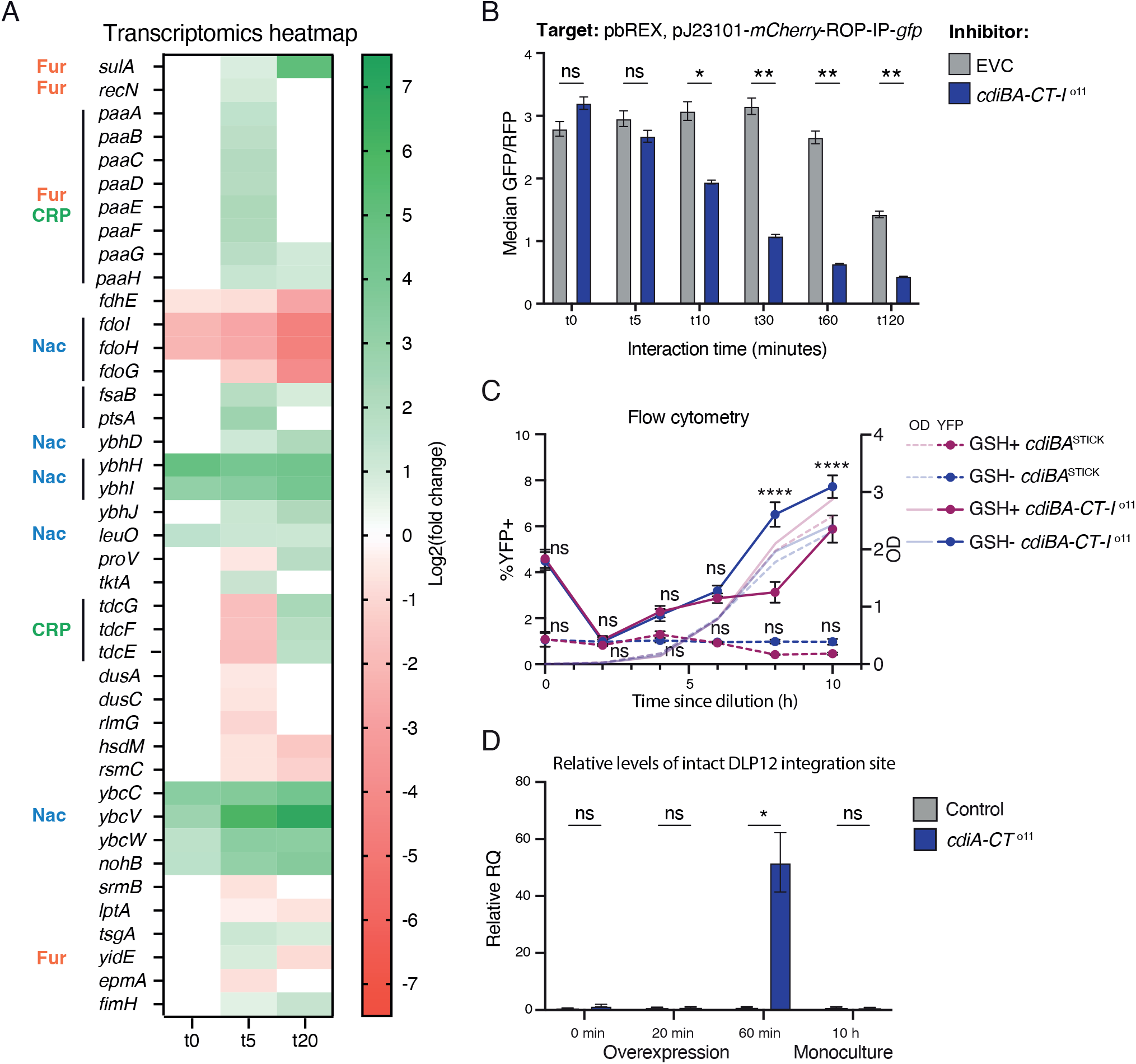
Phenotypic responses to CdiA-CT^o11^ intoxication vary between kin- and non-kin cells. **A)** Heatmap of all genes with altered mRNA levels 5 minutes post-induction of pBAD-*cdiA-CT* ^o11^ as compared to empty vector control (Padj<0.01, N = 3 biological replicates). A subset of known regulators (EcoCyc) is indicated. Coloring indicates Log2fold changes. **B)** Median GFP/RFP level (AU) in targets with the bREX-NADH reporter after 0, 5, 10, 30, 60 or 120 minutes competition against CDI+ (p*cdiBA-CT-I*^o11^, blue) or CDI-(EVC, grey) cells. (N = 3 biological replicates). **C)** Time-resolved enumeration of YFP+ cells in monocultures of MG1655 cells with either p*cdiBA-CT-I* ^o11^ (solid lines) or p*cdiBA*^STICK^ (dashed lines) grown in M9-gly-CAA with (pink) or without (blue) 5 mM glutathione (GSH) for 11 h. (N = 12 biological replicates). Significance between GSH+ and GSH-for the same genotype was determined. **D)** Relative levels of an intact DLP12 attachment site as compared to *purA* ORF, in genomes of *E. coli* MG1655 with pBAD33::*cdiA-CT* ^o11^, pBAD33^empty^ control, *lacA::cdiBA-CT-I* ^o11^ or *lacA::cdiBA* ^STICK^ control. DLP12 excision was quantified using qPCR with primers flanking the integration site where higher presence of an intact locus indicates excision of the prophage. (N = 4 biological replicates). Error bars are SEM. Statistical significance was determined using two-way ANOVA with Tukey’s posthoc test for (C) and Students t-test for (D) where * <0.05, ** <0.01, ***<0,001, ****<0,0001.

Unexpectedly, the majority of the genes with differential expression at 5 minutes are not known to be directly involved in DNA-damage repair. Instead, gene ontology (GO) enrichment analysis using Panther [21] (https://geneontology.org), suggested an over-representation of genes involved in various metabolic processes including phenylacetate catabolism (>40-fold), fatty acid oxidation (>13 fold) and carboxylic acid catabolism (>4-fold) (Table S1). The expression of genes with altered mRNA levels at 5 min is controlled by a diverse set of global transcriptional regulators including Nac, Fur and CRP (Fig. 4A, Table S1)[22], indicating that a slight intoxication results in a global rather than specific changes. Several studies indicate that DNA damaging agents can disturb the redox state of a cell under certain conditions [23, 24]. Redox is known to regulate a wide range of processes in the cell, including the binding ability of the Fur transcription factor [25]. We therefore investigated if CdiA-CT^O11^ activity could affect the redox state of cells by using an established plasmid-based redox sensor, where GFP expression is under the control of the redox sensitive transcription factor Rex [26]. To minimize effects due to handling and change in growth conditions we probed the ratio of NADH+:NAD in targets cells in a competition rather than inducing toxin expression. Already 10 minutes after mixing inhibitors (strain) with sensitive target cells, the ratio of NADH+:NAD dropped significantly (Fig. 4B). No decrease was apparent when competing targets with an empty vector control.

To test if an altered redox state also affects *sulA*-sYFP2 expression in monoculture, we repeated the monoculture setup with or without 5 mM Glutathione (GSH). GSH acts as a redox shunt and has been shown to protect against redox damage by ciprofloxacin [24]. After 8 h, 6% of the cells in a *cdiBA-CT-I*^o11^ monoculture became YFP+ when GSH was absent (GSH-) as compared to 3% when GSH was present (GSH+) (Fig. 4C). After 10h the difference between GSH+/-was less (8% vs 6%), possibly due to GSH consumption (Fig. 4C). Populations with the *cdiBA*^STICK^ showed no increase in YFP+ positive cells in either condition (Fig. 4C). Thus, increasing the reducing capacity of the cell by the addition of GSH mitigates the cellular imbalance in redox state to some extent. Taken together, our results suggest that CdiA-CT^o11^ intoxication results in a shift in redox status, which in turn changes gene-expression of the affected cells.

### Intoxication effects in kin and non-kin cells

A previous study identified a T6SS toxin with deaminase activity, which increased the mutation rate of intoxicated target cells [27]. Theoretically, increasing the mutation rate in a small subpopulation of cells would facilitate adaptation to new niches [28], without affecting the fitness of the entire population [29, 30]. However, none of the genes involved in mutagenic repair (*dinB, umuDC*) were upregulated after 5 min of intoxication, even though 7 to 10-fold induction could be observed after 20min (Fig. S6A). In addition, no increase in mutation rate could be observed in monoculture (Fig. S6BC) or intoxicated cells with immunity that had survived intoxication (Fig. S6DE), suggesting that kin-delivery does not cause subpopulations of hypermutable cells.

DNA damage could also promote phage-mediated horizontal gene transfer (HGT). Prophages are induced by numerous stress signals including the SOS-response [31]. In our dataset, the strongest upregulation occurred in genes encoding prophage proteins. These prophages are unable to form functional phage particles, but over-expression of YbcC and IntD can induce excision of the DLP12 prophage[32]. To assess if intoxication induced DLP12 excision, we used qPCR to detect phage-excission as described previously[32]. Expression of CdiA-CT^o11^ increased DLP12 excision ∼40-fold, suggesting that intoxication of CdiA-CT^o11^ indeed can result in prophage excision. However, we could not observe any prophage excision in monoculture (Fig. 4D), suggesting that, either, kin-delivery does not promote prophage excision or that we are unable to detect such events due to the timing of excision/ putative re-integration.

Taken together, our results suggest that delivery of a CdiA-CT^o11^ results in different outcomes among kin and non-kin cells. Although both effects are dependent on the toxic activity, the level of intoxication seems to guide the response to an altered redox state or to full induction of the DNA damage SOS-response.

## Discussion

Here we show that CDI toxins with DNase activity generate population heterogeneity in terms of gene expression by intoxication of their siblings. This observation agrees with previous reports showing that toxin delivery among siblings, either through CDI or via the type 6 secretion system, can create heterogeneity in gene expression [6, 7]. These reports indicated that toxin delivery among immune kin bacteria induces the RpoS-mediated stress response in some cells [6, 7], but did not untangle whether heterogeneity could arise from i) the delivery event, ii) the toxin - immunity complex acting as transcription factors and/or iii) the toxin activity. The distinction between these mechanisms is important. In i), all toxins delivered would have the same effect whereas for ii) and iii), each toxin could exert different changes. Here we find that CDI-mediated heterogeneity is caused by intoxication (iii), suggesting that each toxin can mediate unique responses. This is supported by the finding that CDI toxins lacking DNase activity are unable to induce *sulA-s*YFP2 (Fig. S7).

Importantly, the changes inflicted in intoxicated cells by the DNase toxin differ between kin-cells and non-immune cells. Although *sulA* expression is induced in both, intoxication of kin-cells results in altered redox status, whereas non-immune cells experience DNA damage, and induce the SOS DNA damage response. These findings are supported by the fact that kin-delivery does not seem to increase mutation rates or promote horizontal gene-transfer (Figs. S6C, E and 4C), whereas at least the latter is observed upon higher intoxication levels (resembling those experienced by non-kin cells). The lack of induction of classical SOS genes 5 minutes after intoxication and in monoculture, despite a clear induction of *sulA*, suggest that *sulA* is regulated by redox as well as LexA, explaining why this reporter responds in monoculture.

What then could be the evolutionary purpose of heterogenicity in redox status in a population? One possibility is that redox-heterogeneity could constitute a form of bet-hedging strategy. Previous findings suggest that CDI mediated population heterogeneity is important for antibiotic tolerance, where some cells are able to survive antibiotic exposure due to growth arrest [7]. In support of this, a recent study links cellular redox status to antibiotic susceptibility in *Pseudomonas* [33]. However, we find that kin-delivery of CdiA-CT^o11^ does not increase antibiotic survival (Fig. S8). In addition, we did not see any increase in mutation rates in response to kin-intoxication. Thus, it does not seem that kin-delivery of toxins is a general bet-hedging strategy. Nonetheless, it is of course possible that this particular toxin enables coping with other types of stresses. More work is needed to shed light on the evolutionary role of kin-delivery of CDI-toxins.

Another possibility is that phenotypic heterogeneity via unequal toxin delivery is important for quasi-multicellular behavior. An important hallmark of this is to acquire heterogeneous responses to the same stimuli among genetically isogenic cells, i.e. polyphenism [34]. Redox status is known to be important for multicellular behavior, regulation of virulence, and cellular responses to environmental cues (reviewed in [35]), suggesting that a heterogeneous redox status could allow for polyphenism in a bacterial population. Thus, although presence of polyphenism itself is not sufficient evidence for division of labor, we suggest that the presence of CDI toxins could lay the basis of multi-cellular behavior, whether this is then used for division of labor or not.

A remaining question is how a DNase toxin changes the redox status of the targeted cell. As also antibiotics with DNA targets, e.g., ciprofloxacin, have previously been shown to change bacterial redox status [23, 24] one might speculate that there is a common pathway through which DNA stress is sensed and forwarded to redox changes in bacteria. Future studies should be aimed at understanding how these processes are intertwined, to get a better understanding of how bacteria respond to various intoxications.

## Methods

### Bacteria and growth conditions

All bacteria used in this study are derivatives of *E. coli* MG1655 and listed in table S1. For media conditions see SI.

### Competition assay

Inhibitor and target strains were grown independently, without antibiotics, to OD_600_∼0.7. The cultures were mixed 5:1 inhibitor:target and grown in 37°C, shaking. Samples were diluted serially and spotted on LB (Fig. S1) or on M9-gly-CAA + L-arabinose (Figs. S3B, 4B and S6) plates with or without KAN for colony counts at every time point. For recovery (Figs 3B, 4B and S6) samples were diluted 1:2 in fresh media containing arabinose for 30 minutes prior to plating and flow cytometry. Competitive index (CI) was calculated as the change in ratio between inhibitor and target over time: CI = (cfu_inhibitor tX_ / cfu_target tX_) / (cfu_inhibitor t0_ /cfu_target t0_). Statistical significance was determined using Students T-test.

### Monoculture experiment

Overnight cultures were diluted 1:1000 into 10 ml fresh medium. Cultures were grown for 11 h with sampling for OD_600_ and flow cytometry measurements at time points -1, 0, 2, 4, 6, 8, 9, 10 and 11 h post-dilution, where -1 h simply indicates pre-dilution. Statistical significance was determined using two-way ANOVA with Tukey’s posthoc test.

### Immunity dilution assay

Strains were grown for approximately 15 h before diluted to OD_600_ ∼0.003 and split into two tubes. Target cells were grown to OD_600_ ∼ 0.5, when L-arabinose was added to one of the tubes. Inhibitor cells were grown for the same time, but with no L-arabinose added. 15 min after L-arabinose addition, all cultures were spun down and resuspended in an equal volume of fresh media. This constituted cycle 0. The culture was diluted 1:5.3 in 20 ml of fresh media and the resulting culture was re-grown for 2 h to OD_600_ ∼0.7. This constituted cycle 1. The same procedure was repeated for a total of 8 cycles with sampling at each cycle for i) OD_600_ measurement (1 ml), ii) a 1:1 competition with the inhibitor for 1 h (3 ml) and iii) western blot (2 ml). The competitive indexes were determined as described above. At cycle 8, the sample was diluted in twice the volume as used for the other cycles (40 ml) and the dilution was split in two flasks. The first flask was treated as all other cycles. The second flask was grown to OD_600_∼0.5, when L-arabinose was added to the culture for 15 min before subjected to the same tests: i) - iii) as the other samples.

## Data Availability Statement

All raw data for the manuscript is available at: https://figshare.com/s/7c41161daa798c92b48a

## Author Contributions

H.E., S.S and S.K conceived the study. H.E. and S.S performed experiments. H.E., S.S, J.K. and S.K analyzed data. H.E., S.S and S.K wrote the manuscript.

## Acknowledgements

We thank Gerhart Wagner for insightful input during the writing of this paper. Sequencing was performed by the SNP&SEQ Technology Platform in Uppsala. The facility is part of the National Genomics Infrastructure (NGI) Sweden and Science for Life Laboratory. The SNP&SEQ Platform is also supported by the Swedish Research Council and the Knut and Alice Wallenberg Foundation.

